# Elevated colonic microbiota-associated paucimannosidic and truncated N-glycans in pediatric ulcerative colitis

**DOI:** 10.1101/2021.05.27.446002

**Authors:** Henghui Li, Xu Zhang, Rui Chen, Kai Cheng, Zhibin Ning, Jianjun Li, Susan Twine, Alain Stintzi, David Mack, Daniel Figeys

## Abstract

Pediatric ulcerative colitis (UC) is a distinct type of inflammatory bowel disease with severe disease activity and rapid progression, which can lead to detrimental life-long consequences. The pathogenesis of pediatric UC remains unclear, although dysbiosis of the gut microbiota has been considered an important factor. In this study, we used mass spectrometry-based glycomic approaches to examine the N-glycans that were associated with the intestinal mucosal-luminal interface microbiota cells of treatment-naïve pediatric UC or control patients. We observed abundant paucimannosidic and other truncated N-glycans that were associated with the microbiota and found that the pediatric UC microbiota samples contained significantly higher levels of these atypical N-glycans compared to those of controls. This study indicates that intestinal N-glycans may be used as novel UC biomarker candidates and the aberrant metabolism of glycans by gut microbiota may be involved in the pathogenesis of UC in children.

---

Ulcerative colitis (UC) is a chronic, continuous inflammatory disease that affects the colon with increasing incidence in both developed and developing countries. The etiology of UC is still unclear, and currently there is no cure for UC in both adults and children. Childhood-onset UC is often more severe than adult UC and approximately 80% of pediatric UC patients have extensive colitis (E3) or pancolitis (E4). In addition, childhood-onset UC progresses much faster than adult-onset UC, and often leads to growth delays and increased risk of long-term comorbidities, such as colorectal cancers ^1^. Therefore, there is an urgent need for early diagnosis and better understanding of the pathogenesis of pediatric UC.

Recent studies highlighted a crucial role of intestinal microbiota in the onset and development of UC in both adult and children ^2^. The intestinal microbiota actively interacts with the host at the mucosal-luminal interface (MLI) by foraging nutrients and regulating immune responses ^3, 4^. The gut microbiota obtains its energy in part by fermenting host-derived and dietary polysaccharides (or glycans) into short-chain fatty acids (SCFAs), which are also known to modulate intestinal homeostasis ^5^. Theses processes are achieved by the presence of extracellular enzymes at the surface of outer membrane or in periplasm of bacteria that breakdown different types of glycans into small oligo- or mono-saccharides. Metagenomic studies have shown that thousands of putative polysaccharide utilization loci (PUL) are present in the gut microbiota, in particular in Bacteroidetes species ^6^.

Recent glycomic analyses of plasma samples revealed alterations of protein glycosylation in both UC and Crohn’s diseases ^7^. Studies of mucin glycans also revealed altered glycan profiles in UC patients ^8^, which might be associated with the aberrant gut microbiota in disease. However, no comprehensive glycomic analysis has been reported for microbiome samples in UC, and it remains unknown whether the dysbiotic gut microbiota in UC leads to altered glycomes in the colon. Therefore, in this study, we performed glycomic analyses of intestinal MLI aspirate samples collected at the ascending colon of UC and non-UC pediatric patients and revealed significant elevation in the relative abundances of paucimannosidic and truncated N-glycans associated with the microbiota. This suggests extensive glycan metabolism by the gut microbiota at the mucosal-luminal interface and point to the potential of using glycans as biomarker for UC or intestinal microbial dysbiosis.

## Results and Discussion

A total of 42 pediatric new-onset UC patients (16 males and 26 females; 13.7 ± 4.0 years [mean ± SD]) and 31 control participants (18 males and 11 females; 13.1 ± 3.5 years [mean ± SD]) were included for analyses in this study (Table 1). Of the UC patients, 30 had pancolitis (Paris classification E4; UC-PCA), while 12 had no-inflammation in the PC (UC-PCN). There were neither significant differences in the sex distribution in each group (Control, P = 0.47; UC, P = 0.16; Binomial two-tailed test) nor significant differences in age (P = 0.25, Mann-Whitney test) between the UC and control participants.

**Table 1.** Clinical characteristics of pediatric UC and control patients

|  | Control (n = 31) | UC (n = 42) |
| --- | --- | --- |
| Male/Female | 18/13 | 16/26 |
| Age [years, median (min-max)] | 15 (3.5-17.7) | 14.2 (5.1-17.5) |
| **Extent** |  |  |
| E1 | NA | 2 |
| E2 | NA | 2 |
| E3 | NA | 8 |
| E4 | NA | 30 |
| **Severity** |  |  |
| S0 | NA | 22 |
| S1 | NA | 20 |
| Severe/moderate/mild | NA | 20/17/5 |

To examine the profiles of glycan present in the intestine of UC or control patients, we first used matrix-assisted laser desorption/ionization-time of flight mass spectrometry (MALDI-TOF-MS) to analyze the MLI aspirate samples collected from ascending colon (PC). This analysis identified 113 N-glycan compositions of which 36 were quantified in >70% samples (Table S1). Interestingly, we observed a general higher peak intensity in the low m/z region (small glycans) in UC compared to control samples (Figure 1A). Five small N-glycan structures were abundantly identified, including three paucimannosidic N-glycans (PMGs; Man_2_GlcNAc_2_Fuc_1_, Man_3_GlcNAc_2_, Man_3_GlcNAc_2_Fuc_1_) and two atypical truncated glycans (Man_4_GlcNAc_2_, Man_2_GlcNAc_3_Fuc_1_). The sum relative intensity of these five rare N-glycans was significantly higher in pancolitis UC (20.9%) than in controls (10.2%; P<0.0001) (Figure 1B). These small atypical N-glycans, including paucimannosidic N-glycan (PMG), have attracted more attentions in recent years and the elevated levels of these unusual glycans has been reported to be associated with many types of cancers, including colorectal cancer ^9^.The abundant expression of these atypical N-glycans in both control and UC MLI microbiota samples (with an observed over-expression in inflamed ascending colon in UC) suggest that the glycans in the gut may be undergoing extensive degradation, which is potentially due to the enzymatic breakdown by gut microbiota given the wide distribution of PULs in metagenome ^6^. These findings also suggest that the aberrant glycan-microbiota interactions might be involved in the development of diseases, such UC and colorectal cancer.

**Figure 1.**
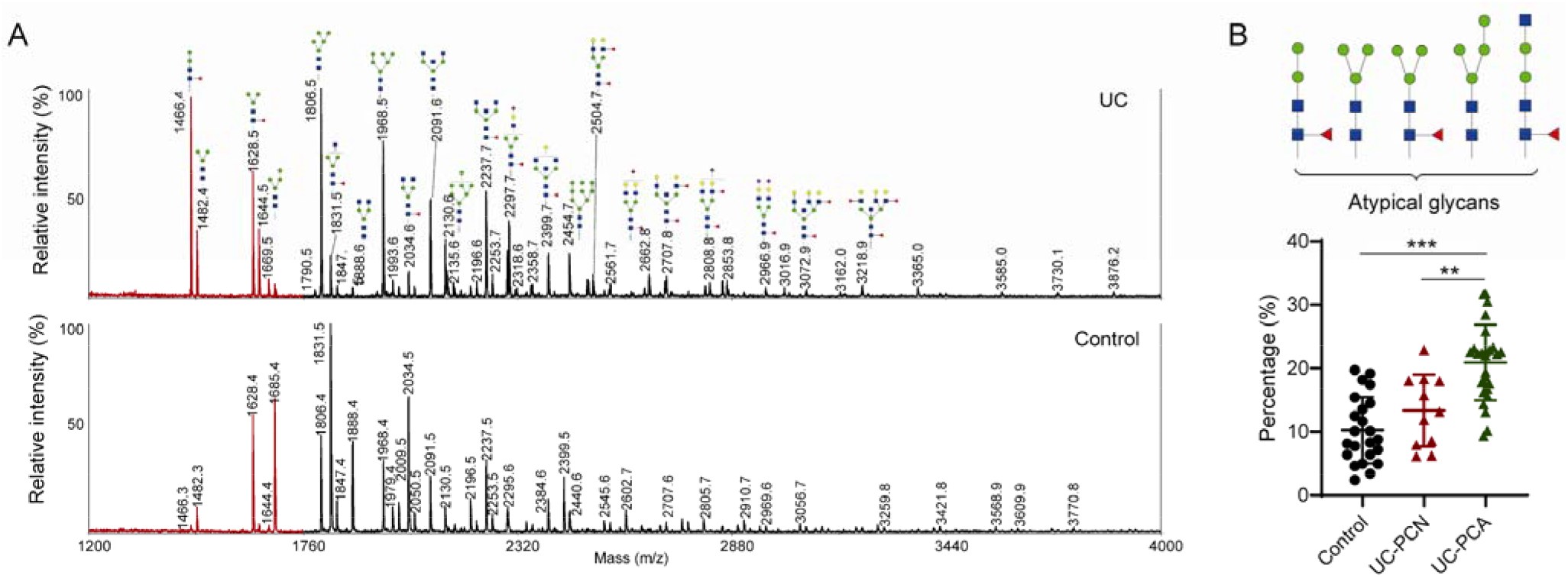
Elevated small N-glycans in MLI microbiome samples of pediatric UC patients. (A) Comparison of patient sample and control samples by MALDI-TOF-MS method. (B) The relative intensity of five atypical structures between patient and control samples by MALDI-TOF-MS method. UC-PCN, non-inflamed ascending colon; UC-PCA, inflamed ascending colon. In the scatter dot plots, individual values (symbol) and mean with standard deviation (SD) are shown. Statistical significance was evaluated using Mann-Whitney U test. **, p < 0.01; ***, p < 0.001.

We then performed partial least squares-discriminant analysis (PLS-DA) using all the 36 glycans quantified across most samples, which successfully discriminated UC from controls with an accuracy of 0.91 (Q^2^=0.43, R^2^=0.57). By using a variable importance of projection (VIP) value threshold of 1.0, eleven glycans were identified as differentially abundant in UC compared to controls, and two (Man_2_GlcNAc_2_Fuc_1_ and Man_3_GlcNAc_3_Fuc_1_Gal_1_NeuNAc_1_) of which had an area under the receiver operating characteristic curve (AUC) value of 0.9 and higher (Figure 2 and Table S1), representing promising biomarker candidates for pediatric UC.

**Figure 2.**
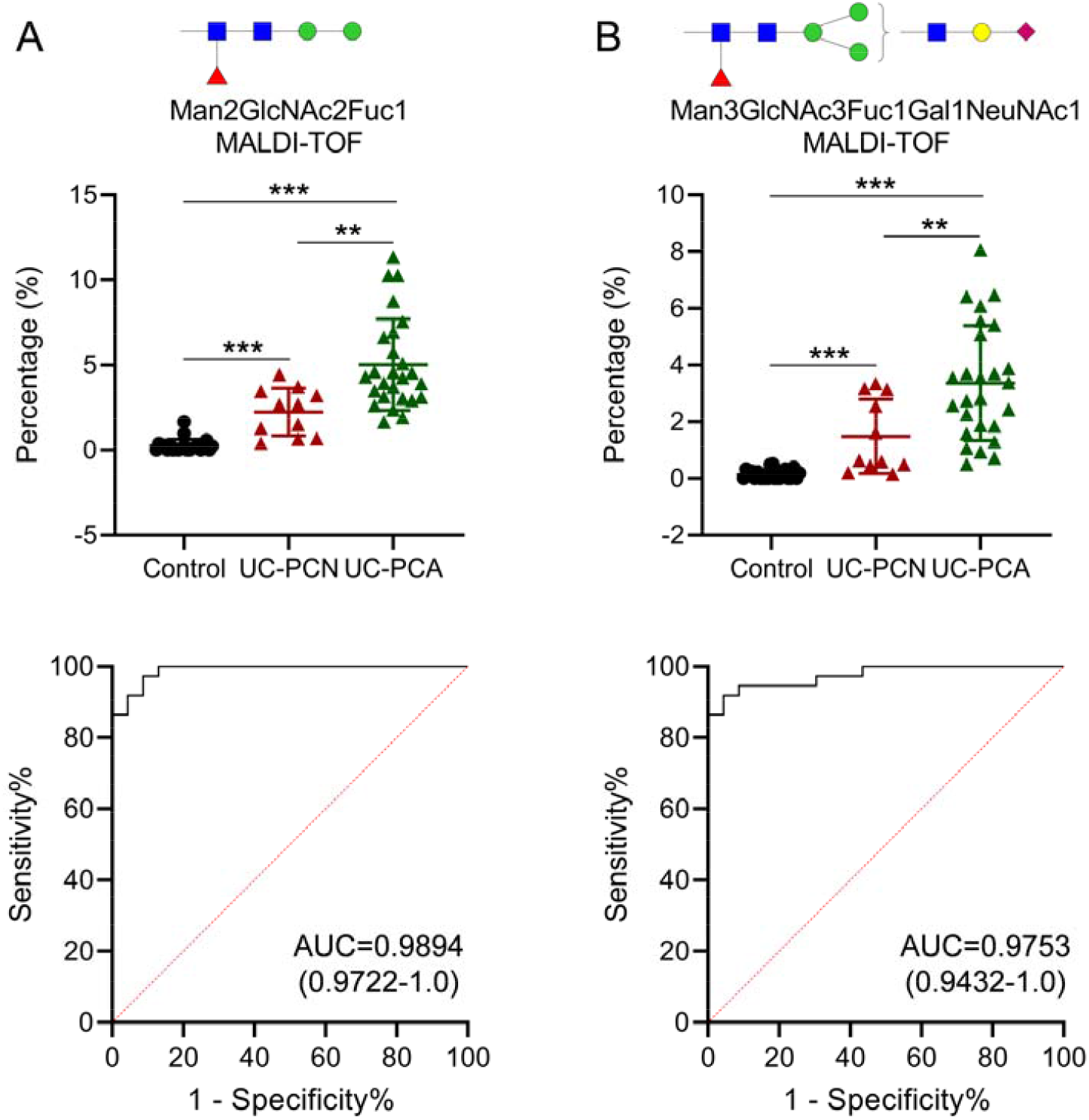
Relative expression levels of two differentially abundant N-glycans as measured by MALDI-TOF-MS. (A-B) Relative expression levels (upper panel) and ROC curve (lower panel) were shown for each differentially abundant N-glycan. Glycan compositions are indicated at the top. AUC and 95% confidence interval are indicated. In the scatter dot plots, individual values (symbol) and mean with standard deviation (SD) are shown. Statistical significance was evaluated using Mann-Whitney U test. **, p < 0.01; ***, p < 0.001.

To perform a more comprehensive glycomic analysis, we then applied porous graphitized carbon-based liquid chromatography-tandem mass spectrometry (PGC-LC-MS) to analyze the MLI samples collected from pediatric patients, which successfully quantified 127 glycans with valid values in >90% samples. Unsupervised principal component analysis (PCA) using all the 127 quantified glycans revealed obvious differences in glycan profiles between UC and controls (Figure 3). PLS-DA efficiently discriminate UC from controls with an accuracy of 0.91 (Q^2^=0.67, R^2^=0.70). Among the nine glycans with VIP value of >1.0, four showed an AUC > 0.9 and all of them are paucimannosidic or truncated glycans (Figure 4A-D and Table S1).

**Figure 3.**
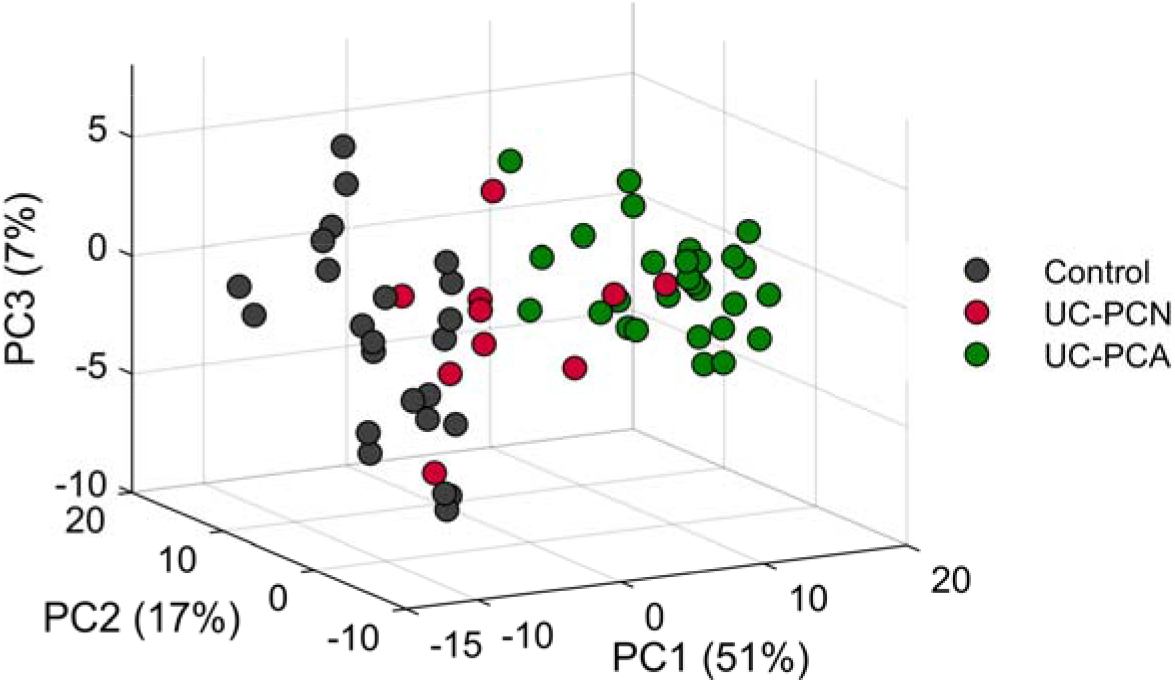
PCA score plot of quantified microbiota-associated N-glycans in pediatric UC.

**Figure 4.**
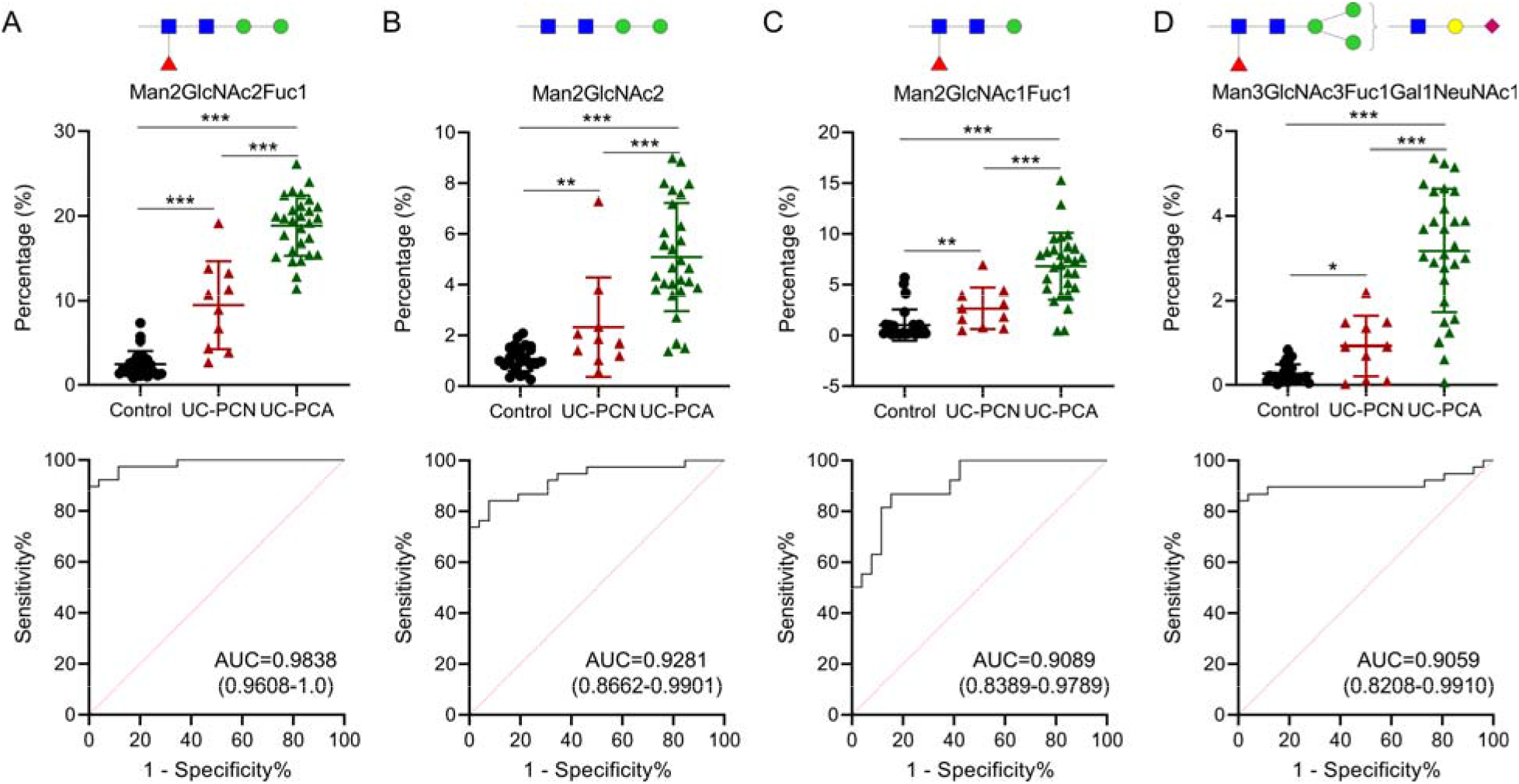
Relative expression levels of four differentially abundant N-glycans as measured by PGC-LC-MS. (A-D) Relative expression levels (upper panel) and ROC curve (lower panel) were shown for each differentially abundant N-glycan. Glycan compositions are indicated at the top. AUC and 95% confidence interval are indicated. In the scatter dot plots, individual values (symbol) and mean with standard deviation (SD) are shown. Statistical significance was evaluated using Mann-Whitney U test. *, p < 0.05; **, p < 0.01; ***, p < 0.001.

Man_2_GlcNAc_2_Fuc_1_ had the highest AUC (0.9838, 95% CI: 0.9608–1.0) with a relative abundance of 2.49% in controls, 10.75% in UC-PCN and 18.39% in UC-PCA, representing the most abundant glycan in MLI aspirate samples. Interestingly, Man_2_GlcNAc_2_Fuc_1_ was also identified as differentially abundant glycan in MALDI-TOF-MS data set and had the highest AUC (0.9894, 95% CI: 0.9722–1.0) (Figure 2A). Man_3_GlcNAc_3_Fuc_1_Gal_1_NeuNAc_1_, a biantennary glycan, was the other glycan structure that were identified as differentially abundant glycans in UC compared to controls in both PGC-LC-MS (AUC, 0.9059, 95% CI: 0.8208-0.99) and MALDI-TOF-MS data sets (AUC, 0.9753, 95% CI: 0.9432-1.0). Correlation analyses of their relative abundances quantified using the two MS approaches showed high accordance for both Man_2_GlcNAc_2_Fuc_1_ (Spearman’s r = 0.75, p < 0.0001) and Man_3_GlcNAc_3_Gal_1_NeuNAc_1_ (Spearman’s r = 0.82, p < 0.0001) (Figure 5), suggesting high data quality and validity of these identified alterations of glycan composition at the intestinal mucosal-luminal interface of pediatric UC patients.

**Figure 5.**
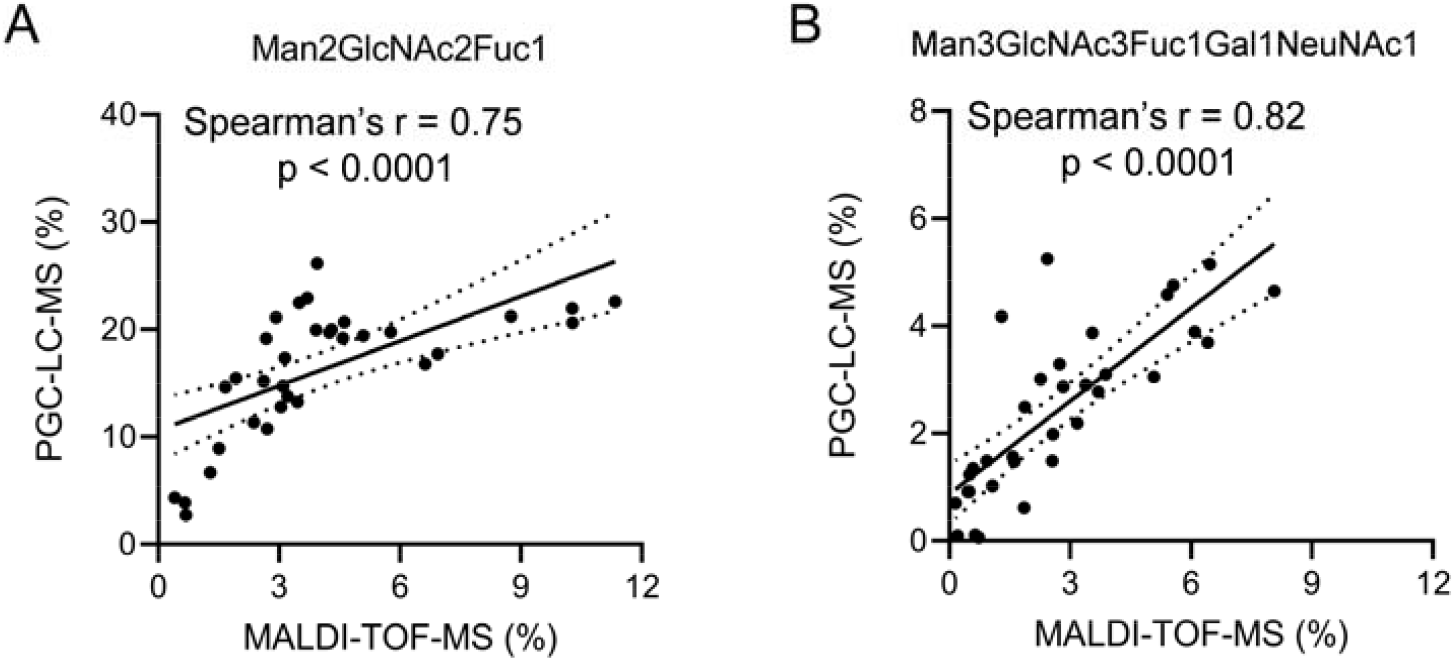
Correlation of the relative expression levels of N-glycans quantified using PGC-LC-MS and MALDI-TOF-MS methods. (A) Man_2_GlcNAc_2_Fuc_1_. (B) Man_3_GlcNAc_3_Gal_1_NeuNAc_1_. Spearman’s correlation coefficient r and p are indicated.

As mentioned above, the intestinal microbiota has developed diverse glycan-degrading strategies, and the microbial genes for glycan degradation (i.e., PULs) are mostly present in species from Bacteroidetes ^6^ and some butyrate-producing Firmicutes species ^10^. *Bacteroides thetaiotaomicron*, an abundant *Bacteroides* species in human gut, encodes complex and diverse genetic apparatus, enabling the access of various structural variants of complex N-glycans that might be present in the gut ^11^. In UC patients, a decreased level of SCFA-producing species in Firmicutes and decreased species diversity within Bacteroidetes were previously reported ^2, 12^. By using metaproteomic approach, we also previously reported that the MLI microbiota in pediatric UC patients displayed higher relative abundance of *Bacteroides* and lower abundance of *Feacalibacterium*, an abundant SCFA-producer in human gut, compared to those of control patients ^4^. The dysbiotic gut microbiota in UC patients may thereby lead to aberrant capability and/or specificity to breakdown polysaccharides into small glycans, such as PMGs. It remains unclear which microbial species or metabolic pathway changes lead to the elevated levels of these truncated glycans in the intestine of UC patients, however, the findings in the current study suggest that the aberrant glycan utilization of gut microbes might be involved in the onset and development of UC.

To conclude, we demonstrated that paucimannosidic and other truncated N-glycan structures were abundantly expressed in both control and UC MLI aspirate samples, in particular the inflamed colon aspirates in UC patients, by glycomic approach. These alterations of glycan composition in the intestine of UC patients might be due to the dysbiotic microbiota that can lead to aberrant capability and/or specificity to breakdown polysaccharides into small glycans. Our preliminary findings indicate that the intestinal glycans, including the truncated ones, may serve as a source of promising biomarker candidates for the diagnosis of UC although further mechanistic studies and validation in larger patient cohorts are needed.

## Methods

### Patients, MLI sample collection and processing

Pediatric patients (<18 years old) that underwent diagnostic colonoscopy for suspected UC from November 2014 to November of 2019 at CHEO were considered and a total of 42 pediatric new onset UC patients (16 males and 26 females; 13.7 ± 4.0 years [mean ± SD]) and 31 control participants (18 males and 11 females; 13.1 ± 3.5 years [mean ± SD]) were included for this study. The study was approved by the Research Ethic Board of the Children Hospital of Eastern Ontario (CHEO), Ottawa, ON, Canada. The exclusion and inclusion criteria were described previously ^4, 13, 14^. UC diagnosis and classification were performed according to the standard criteria as described previously ^15^, and patients with visually and histologically normal mucosa were considered as control. UC disease activity was evaluated using PUCAI (Pediatric Ulcerative Colitis Activity Index) ^16^. All patient information was recorded in Redcap that was hosted in CHEO (https://redcap.cheori.org/).

MLI aspirate samples were collected from the ascending colon during colonoscopy following a standard procedure ^17^. Briefly, during colonoscopy, any loose fluid and debris was aspirated away, and then sterile water was flushed onto the mucosa and the resulting mixture was collected and kept on ice for processing immediately. Any debris was first removed by centrifugation at 700 g for 5 min and the resulting supernatant was subjected for centrifugation at 14,000□g, 4□°C for 20□min to harvest microbial cells. The latter was washed once with ice-cold PBS and stored in -80□°C for further processing.

Protein extraction was performed using lysis buffer containing 4 % (w/v) sodium dodecyl sulfate, 8□M urea, 50□mM Tris-HCl, and protease inhibitor as described previously ^18^. Three ultrasonications (30□s each with 30s interval on ice) were applied to facilitate microbial cell disruption. Cell debris was removed through centrifugation and the resulting supernatant was precipitated using acidified acetone/ethanol buffer at −20□°C overnight. Precipitated proteins were then washed three times with ice-cold acetone for further glycomic analysis.

### MALDI-TOF-MS analysis

100 ug of proteins were dissolved in 50 μL HEPES (50mM, pH8.0) containing 1% RapiGest SF surfactant, 50 mM dithiothreitol (DTT). The proteins were denatured at 95 °C for 5 min, and cooled down to 50 °C for releasing reaction with the addition of 1.5 μL Rapid PNGase F. N-Succinimidyloxycarbonylmethyl tris(2,4,6-trimethoxyphenyl)-phosphonium bromide (TMPP-Ac-OSu) labeling and purification were then performed according to previous study ^19^. Briefly, a freshly prepared 50 μL 0.02mg/μL TMPP-Ac-OSu in acetonitrile was added to the sample with vigorously agitation for 30 min at room temperature. 500 μL binding solution was then added for loading to an hydrophilic interaction liquid chromatography (HILIC) method with home-packed microcrystalline cellulose solid-phase extraction (SPE) column ^20^. The derivatized glycans were eluted by 1 mL of ethanol/H2O (1:1, v/v) and dried by a SpeedVAC concentrator for further derivatization using methylamine as previously described ^21^. Briefly, the glycan was dissolved in 50 μL of dimethylsulfoxide containing 1 M methylamine hydrochloride, 0.5 M N-methylmorpholinerior, and 50 mM PyAOP (Cat851221, Sigma-Aldrich) and incubated at room temperature for 30 min. Glycan purification was performed as described above using HILIC SPE and the eluted derivatized glycan was used for MALDI-TOF-MS measurement.

MALDI-TOF-MS was performed using 4800 MALDI-TOF/TOF mass spectrometer (AB SCIEX) equipped with a 355 nm Nd: YAG laser in the reflector positive mode. Derivatized glycan samples were mixed with 10 mg/ml dihydroxybenzoic acid matrix, loaded onto the stainless steel MALDI plate, and dried at room temperature. A total of 1000 laser shots were employed in each sample spot. The glycan annotation was performed according to the consortium for functional glycomics and the cartoons of possible structures of glycans were adapted from Glycoworkbench ^22^. The spectra were exported as the text files by Data Explorer 4.0 (AB SCIEX) for further analysis. Normalization was performed by first summing the intensities of all glycans of interest as 100% and calculating the relative abundance of each signal associated with each ion ^23^. The relative abundance of each ion was then used for further data analysis.

### PGC nano-LC MS analysis

100ug proteins were re-suspended in 100μL 20 mM PBS buffer (pH= 7.5) with 0.1% SDS and 5 mM DTT. Following denaturing at 95 °C for 5 min and cooling to room temperature, 5 μL of 10% octylphenoxypolyethoxyethanol (NP-40) was added and the mixture was incubated with 5 units of PNGase F at 37 °C overnight. The samples were then dried and purified using 96-well hypercarb PGC plates (Thermo fisher) according to the manufacture’s instruction. The glycans were eluted with 1 mL 0.5% TFA/40% acetonitrile and dried using SpeedVAC. The resulting N-glycan was then derivatized by methylamine and purified using HILIC SPE as described above. The glycan was analyzed by an Eksigent nano-LC that was connected in-line with an Q-Exactive HFX mass spectrometer ^24^. The separation of glycan was performed on an analytical column packed with PGC phase beads (Hypercarb 3 µm; pore size 250 Å; Thermo Fisher Scientific) with 30 min from 5 to 35% acetonitrile (v/v) at a flow rate of 200 nl/min. The full scan MS were acquired over range 300-3500 (m/z) with the mass resolution setting 70000 at m/z 400. Maximum injection time 100 ms; AGC target value of 1e6. The 12 most intense ions were selected for MSMS detection with stepped collision energy (15, 20, and 25), exclusion ion charge ≥5; resolution 17500, AGC target 1e5, and maximum injection time 120 ms.

MS raw data were exported as mzXML files. The N-linked glycan composition was identified and verified using both MSMS with a 10ppm mass tolerance and retention-time. MS/MS spectra were inspected for each structure based on the feature ions. Glycans were relatively quantified to the total intensity by integration of the extracted ion chromatogram peak area (EIC). Normalization was performed by the sum of the chromatographic peak areas for all charge states of the observed N-linked glycans.

### Statistical data analysis

The glycan compositions that were quantified in >70% and >90% samples for MALDI-TOF-MS and PGC-LC-MS data set, respectively, were used for further biomarker analysis. PLS-DA and ROC curve analysis were performed using MetaboAnalyst 4.0 ^25^. PCA analysis was performed in MATLAB (version 2019a, The MathWorks, Inc). Statistical analysis and figure plotting were performed in Prism (version 8, GraphPad).

## Acknowledgments

This work was supported by funding from the Natural Sciences and Engineering Research Council of Canada (NSERC), the Government of Canada through Genome Canada and the Ontario Genomics Institute (OGI-114 & OGI-149), CIHR grant number GPH-129340 and MOP-114872, and the Ontario Ministry of Economic Development (Project number 13440). DF acknowledges a Distinguished Research Chair from the University of Ottawa. We acknowledge Ruth Singleton at the Children’s Hospital of Eastern Ontario, Mila Tepliakova and James Butcher at the University of Ottawa for their help in collecting and processing intestinal aspirate samples.

## Author information

### Contributions

D.F., A.S., and D.M. designed the study. D.M. collected patient samples and clinical data. H.L., X.Z., R.C., Z.N. and K.C. performed the experiments and data analysis. H.L., X.Z., and D.F. wrote the manuscript and all authors participated in the data interpretation, discussion, and edits of the manuscript.

**Corresponding authors**

Correspondence to Daniel Figeys.

## Ethics declarations

### Conflict of interest

D.F., D.M. and A.S. have co-founded MedBiome, a clinical microbiomics company. The remaining authors declare no competing interests.

## References

1. Sauer CG, Kugathasan S. Pediatric inflammatory bowel disease: highlighting pediatric differences in IBD. Gastroenterology clinics of North America 38, 611–628 (2009).

2. Lloyd-Price J, et al. Multi-omics of the gut microbial ecosystem in inflammatory bowel diseases. Nature 569, 655–662 (2019).

3. Sonnenburg JL, et al. Glycan foraging in vivo by an intestine-adapted bacterial symbiont. Science 307, 1955–1959 (2005).

4. Zhang X, et al. Metaproteomics reveals associations between microbiome and intestinal extracellular vesicle proteins in pediatric inflammatory bowel disease. Nature communications 9, 2873 (2018).

5. Hill MJ. Bacterial fermentation of complex carbohydrate in the human colon. European journal of cancer prevention : the official journal of the European Cancer Prevention Organisation 4, 353–358 (1995).

6. Lapebie P, Lombard V, Drula E, Terrapon N, Henrissat B. Bacteroidetes use thousands of enzyme combinations to break down glycans. Nature communications 10, 2043 (2019).

7. Clerc F, et al. Plasma N-Glycan Signatures Are Associated With Features of Inflammatory Bowel Diseases. Gastroenterology 155, 829–843 (2018).

8. Lennon G, et al. Influences of the colonic microbiome on the mucous gel layer in ulcerative colitis. Gut microbes 5, 277–285 (2014).

9. Chatterjee S, et al. Protein Paucimannosylation Is an Enriched N-Glycosylation Signature of Human Cancers. Proteomics 19, e1900010 (2019).

10. La Rosa SL, et al. The human gut Firmicute Roseburia intestinalis is a primary degrader of dietary beta-mannans. Nature communications 10, 905 (2019).

11. Briliute J, et al. Complex N-glycan breakdown by gut Bacteroides involves an extensive enzymatic apparatus encoded by multiple co-regulated genetic loci. Nature microbiology 4, 1571–1581 (2019).

12. Ishikawa D, et al. The Microbial Composition of Bacteroidetes Species in Ulcerative Colitis Is Effectively Improved by Combination Therapy With Fecal Microbiota Transplantation and Antibiotics. Inflammatory bowel diseases 24, 2590–2598 (2018).

13. Starr AE, et al. Proteomic analysis of ascending colon biopsies from a paediatric inflammatory bowel disease inception cohort identifies protein biomarkers that differentiate Crohn’s disease from UC. Gut 66, 1573–1583 (2017).

14. Deeke SA, et al. Mucosal-luminal interface proteomics reveals biomarkers of pediatric inflammatory bowel disease-associated colitis. The American journal of gastroenterology 113, 713–724 (2018).

15. Bousvaros A, et al. Differentiating ulcerative colitis from Crohn disease in children and young adults: report of a working group of the North American Society for Pediatric Gastroenterology, Hepatology, and Nutrition and the Crohn’s and Colitis Foundation of America. Journal of pediatric gastroenterology and nutrition 44, 653–674 (2007).

16. Turner D, et al. Development, validation, and evaluation of a pediatric ulcerative colitis activity index: a prospective multicenter study. Gastroenterology 133, 423–432 (2007).

17. Jimenez-Rivera C, Haas D, Boland M, Barkey JL, Mack DR. Comparison of two common outpatient preparations for colonoscopy in children and youth. Gastroenterology research and practice 2009, 518932 (2009).

18. Zhang X, Li L, Mayne J, Ning Z, Stintzi A, Figeys D. Assessing the impact of protein extraction methods for human gut metaproteomics. J. Proteomics 180, 120–127 (2018).

19. Gao W, et al. Rapid and sensitive analysis of N-glycans by MALDI-MS using permanent charge derivatization and methylamidation. Talanta 161, 554–559 (2016).

20. Zhang Q, Li H, Feng X, Liu BF, Liu X. Purification of derivatized oligosaccharides by solid phase extraction for glycomic analysis. PloS one 9, e94232 (2014).

21. Liu X, Qiu H, Lee RK, Chen W, Li J. Methylamidation for NeuNAcloglycomics by MALDI-MS: a facile derivatization strategy for both alpha2,3-and alpha2,6-linked NeuNAclic acids. Analytical chemistry 82, 8300–8306 (2010).

22. Ceroni A, Maass K, Geyer H, Geyer R, Dell A, Haslam SM. GlycoWorkbench: a tool for the computer-assisted annotation of mass spectra of glycans. Journal of proteome research 7, 1650–1659 (2008).

23. Kang P, Madera M, Alley WR, Jr., Goldman R, Mechref Y, Novotny MV. Glycomic Alterations in the Highly-abundant and Lesser-abundant Blood Serum Protein Fractions for Patients Diagnosed with Hepatocellular Carcinoma. International journal of mass spectrometry 305, 185–198 (2011).

24. Li H, et al. Chemoenzymatic Method for Glycoproteomic N-Glycan Type Quantitation. Analytical chemistry 92, 1618–1627 (2020).

25. Chong J, et al. MetaboAnalyst 4.0: towards more transparent and integrative metabolomics analysis. Nucleic acids research 46, W486–W494 (2018).

